# Mutations that adapt SARS-CoV-2 to mustelid hosts do not increase fitness in the human airway

**DOI:** 10.1101/2021.08.20.456972

**Authors:** Jie Zhou, Thomas P. Peacock, Jonathan C. Brown, Daniel H. Goldhill, Ahmed M.E. Elrefaey, Rebekah Penrice-Randal, Vanessa M. Cowton, Giuditta De Lorenzo, Wilhelm Furnon, William T. Harvey, Ruthiran Kugathasan, Rebecca Frise, Laury Baillon, Ria Lassaunière, Nazia Thakur, Giulia Gallo, Hannah Goldswain, I’ah Donovan-Banfield, Xiaofeng Dong, Nadine P. Randle, Fiachra Sweeney, Martha C. Glynn, Jessica L. Quantrill, Paul F. McKay, Arvind H. Patel, Massimo Palmarini, Julian A. Hiscox, Dalan Bailey, Wendy S. Barclay

**Affiliations:** Department of Infectious Disease, Imperial College London, UK; The Pirbright Institute, Woking, Surrey, UK; Institute of Infection, Veterinary and Ecology Sciences, University of Liverpool, UK; MRC-University of Glasgow Centre for Virus Research, Glasgow, UK; Virus & Microbiological Special Diagnostics, Statens Serum Institut, Copenhagen, Denmark; The Jenner Institute, Nuffield Department of Medicine, University of Oxford, UK; Infectious Diseases Horizontal Technology Centre (ID HTC), A*STAR, Singapore

**Keywords:** SARS-CoV-2, COVID-19, coronavirus, mink, ferret, antigenicity, ACE2

## Abstract

SARS-CoV-2 has a broad mammalian species tropism infecting humans, cats, dogs and farmed mink. Since the start of the 2019 pandemic several reverse zoonotic outbreaks of SARS-CoV-2 have occurred in mink, one of which reinfected humans and caused a cluster of infections in Denmark. Here we investigate the molecular basis of mink and ferret adaptation and demonstrate the spike mutations Y453F, F486L, and N501T all specifically adapt SARS-CoV-2 to use mustelid ACE2. Furthermore, we risk assess these mutations and conclude mink-adapted viruses are unlikely to pose an increased threat to humans, as Y453F attenuates the virus replication in human cells and all 3 mink-adaptations have minimal antigenic impact. Finally, we show that certain SARS-CoV-2 variants emerging from circulation in humans may naturally have a greater propensity to infect mustelid hosts and therefore these species should continue to be surveyed for reverse zoonotic infections.

## Introduction

SARS-CoV-2 is a betacoronavirus that is thought to have emerged from an animal source in 2019 and rapidly spread by human-to-human transmission across the globe causing the COVID-19 pandemic. SARS-CoV-2 is transmitted efficiently by the airborne route due to its ability to efficiently enter cells in the upper respiratory tract. The spike glycoprotein is responsible for host receptor binding and membrane fusion of coronaviruses. SARS-CoV-2 spike binds to host angiotensin-converting enzyme 2 (ACE2) via the receptor binding domain (RBD) and is activated by TMPRSS2 protease expressed at the apical surface of the airway epithelium to mediate fusion ^1^. In addition, compared to closely related coronaviruses, SARS-CoV-2 spike contains a tract of basic amino acids at the S1/S2 cleavage site that can be recognised by furin, enabling spike to be efficiently primed for fusion by TMPRSS2. This allows rapid fusion of spike at the cell surface and avoids restriction factors present in the late endosome and endolysosome ^2,3^.

A series of molecular interactions between amino acids in the spike RBD and the interacting surface of ACE2 result in SARS-CoV-2 binding to human ACE2 with high affinity ^4,5^. SARS-CoV-2 shows a broad host tropism and can experimentally infect many animal species, largely determined by the efficiency with which spike can interact with the animal ACE2 orthologues ^5,6^. For example, mice are inefficiently infected by early SARS-CoV-2 isolates unless they are engineered to transgenically express human ACE2 or SARS-CoV-2 is adapted to murine ACE2 by serial mouse passage ^7,8^.

From April 2020, reverse-zoonotic outbreaks (i.e. transmitted from humans into animals; also known as zooanthroponosis) of SARS-CoV-2 in mink farms were reported in the Netherlands, the USA, France, Spain, Denmark, Italy, Sweden, Canada, Greece, Lithuania, and Poland ^9–13^. Multiple reverse zoonotic events introduced the virus from farm workers into densely populated farms that then supported rapid transmission between animals ^12,13^. Sequence analyses revealed several mutations in spike enriched after circulation in mink, most commonly the amino acid substitution Y453F or N501T; residues that map to the RBD of spike protein ^11–13^. From June to November 2020, an outbreak of SARS-CoV-2 infections occurred among farmed mink in Denmark, with continuous spillover to farm workers and local communities with viruses harbouring Y453F ^13,14^. This large-scale outbreak resulted in an estimated 4000 mink-associated human cases and promoted the Danish government to cull all 17 million farmed mink in the country and several countries imposed total travel bans on the affected regions ^15^. Of particular concern was the increased acquisition of spike mutations in mink-associated SARS-CoV-2 viruses, as demonstrated by the Cluster 5 variant identified in September 2020 which had several additional changes in the spike glycoprotein including, Δ69-70 in the N-terminal domain (NTD), and I692V and M1229I in S2 ^16^. Early data indicated a possible change in antigenicity whereby Cluster 5 variant virus might be less readily neutralised by antibodies in convalescent sera from individuals infected by earlier variants ^16,17^.

Ferrets are closely related to mink (both belong to the family *Mustelidae*) and have been used extensively as models for transmission of influenza virus due to their high susceptibility, comparable tissue tropism, and clinical signs similar to those seen in infected humans ^18^. Consequently, ferrets have also been extensively characterised as models for SARS-CoV-2 transmission experiments and can support infection and transmission ^3,19,20^. During experimental infection of SARS-CoV-2 in ferrets, several groups have independently reported mink-associated spike mutations Y453F or N501T ^12,19,21^. Mink and ferret ACE2 are extremely similar with no amino acid differences that map to the ACE2/SARS-CoV-2 Spike interface (Supplementary Figure 1). Interestingly, both the Y453F and the N501T substitutions have also been associated with increased binding to human ACE2 ^22^. It is possible these mutations may act to non-specifically increase binding to several groups of mammalian ACE2 proteins. A similar mutation N501Y, which is also associated with increased human ACE2 binding is often found upon mouse adaptation of SARS-CoV-2 and is present in several human SARS-CoV-2 variants of concern showing signs of higher transmissibility ^23–25^. It has been hypothesised for influenza virus that increasing receptor binding avidity can result in non-specific antibody escape as the viral glycoprotein-host receptor interaction begins to outcompete that of antibody/viral glycoprotein ^26^.

In this study, we risk assess mink or ferret adapted viruses and mutations to determine the threat that viruses adapted to mustelid species could pose to humans, and what impact they could have on global vaccine efforts.

## Results

### Y435F and N501T substitutions in the spike are detected in viruses transmitted between ferrets

In a previous study, we experimentally infected four donor ferrets and tested the ability of SARS-CoV-2 to transmit to individually co-housed naive animals ^3^. The early wildtype (WT) virus isolate, England/2/2020, which contains 614D, transmitted efficiently to 2 of 4 ferrets in direct contact with two infected donor animals (donor #1 and #2; Figure 1A). Here, we sequenced the spike RBD of virus extracted from nasal wash obtained at early and late time point from the direct contact ferrets. Of the two transmitted virus isolates, at the consensus level, one had gained N501T in the spike protein while the other had a mixture of Y453F and N501T. Both Y453F and N501T have previously been associated with experimental ferret adaptation of SARS-CoV-2 ^19,21^. To investigate the dynamics of ferret adaptation in more detail, virus samples were deep sequenced from various times point across the 2 successful ferret transmission chains (donor #1 to contact #1 and donor #2 to contact #2; Figure 1A). In donor animal #1, the virus rapidly gained majority N501T with Y453F as a minor variant. However, by day 5, both mutations were present in equal amounts with no detectable WT spike. In the matched contact animal (contact #1), the transmitted virus population included a mixture of Y453F with a minority of N501T and Y453F continued to predominate between days 4-6. In Donor/Contact pair #2, again both mutations were detected but N501T predominated across both animals at all time points tested. N501T alone predominated in the day 2 nasal washes from the two donor animals that did not transmit to their direct contacts (98% in one, 94% in the other), with remaining reads showing WT spike ^3^. In the initial virus inoculum, N501T was detected at levels below 1% of total reads while Y453F was not detected at all (read depth ≈7000). No other consensus level mutations arose in any of the donor or contact ferrets anywhere else in the genome.

**Figure 1.**
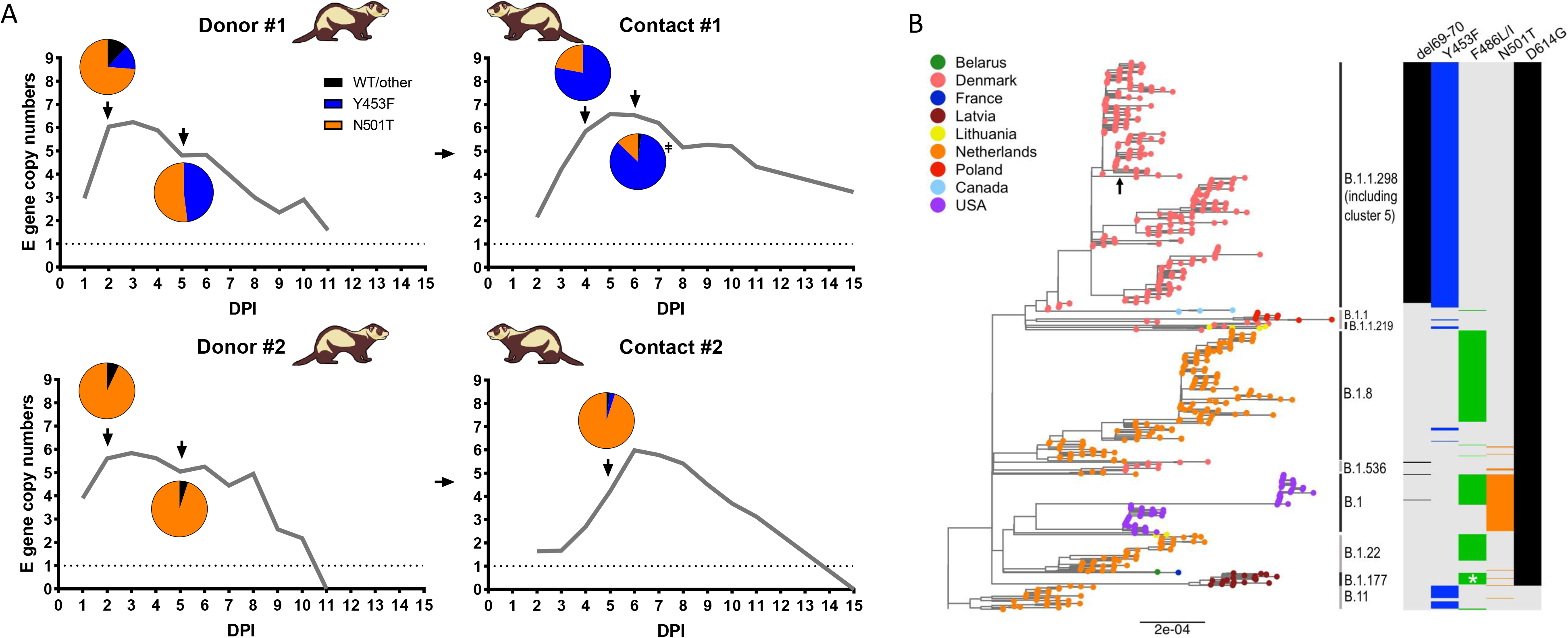
Passage of SARS-CoV-2 in ferrets results in spontaneous emergence of the mink-associated mutations Y453F and N501T. A) Ferret transmission chains from a previous study were deep sequenced to investigate any changes that occurred during infection and transmission of ferrets. Grey lines indicate previously described RNA shedding patterns seen in each ferret – pie charts indicate RBD mutations seen at each time point (as determined by deep sequencing) indicated by a black arrow. B) Maximum-likelihood phylogeny of SARS-CoV-2 genomes sampled from American mink (*Neogale vison*, formerly *Neovison vison*), highlighting the spike mutations del69-70, Y453F, F486L or F486I, N501T, and D614G. Tip nodes are shown as points coloured by sampling location, according to the colour key. Columns to the right show the presence of either the wild-type amino acid(s) (light grey) or the mutations annotated above (coloured bars). Major epidemiological lineages designated with the Pango nomenclature system are labelled. Black arrow indicates the branch that constitutes the Danish mink strain known as cluster 5. At position 486, mutant viruses possessed 486L (leucine) except for a monophyletic clade formed of 20 sequences sampled in Latvia that possessed 486I (isoleucine) that are marked by a white asterisk.

Interesting, by investigating all SARS-CoV-2 sequences isolated from mink reported on GISAID, we and others noted that N501T, Y453F, as well as F486L have independently arisen multiple times in mink, and in multiple lineages as illustrated in Figure 1B ^11,12^. These observations further imply that these mutations are strongly associated with mustelid adaptation (Figure 1B).

Of all the mink-adapting substitutions, Y453F has been more frequently associated with spillback from mink into humans, including Cluster 5 in Denmark. To further investigate the effects of the Y453F substitution, we isolated virus from contact #1 from day 6 in Vero cells (‘Ferret P2’) and validated that the sequence change was maintained in the titrated virus stock (Figure 2A). The Vero grown virus stock was, in the majority, Y453F (^∼^96%) with very minor variants, N501T and WT RBD also present (<5%). Outside of spike, the virus contained an additional mutation, S6L in the envelope gene (E) which was present in >70% of reads in the Vero grown stock.

**Figure 2.**
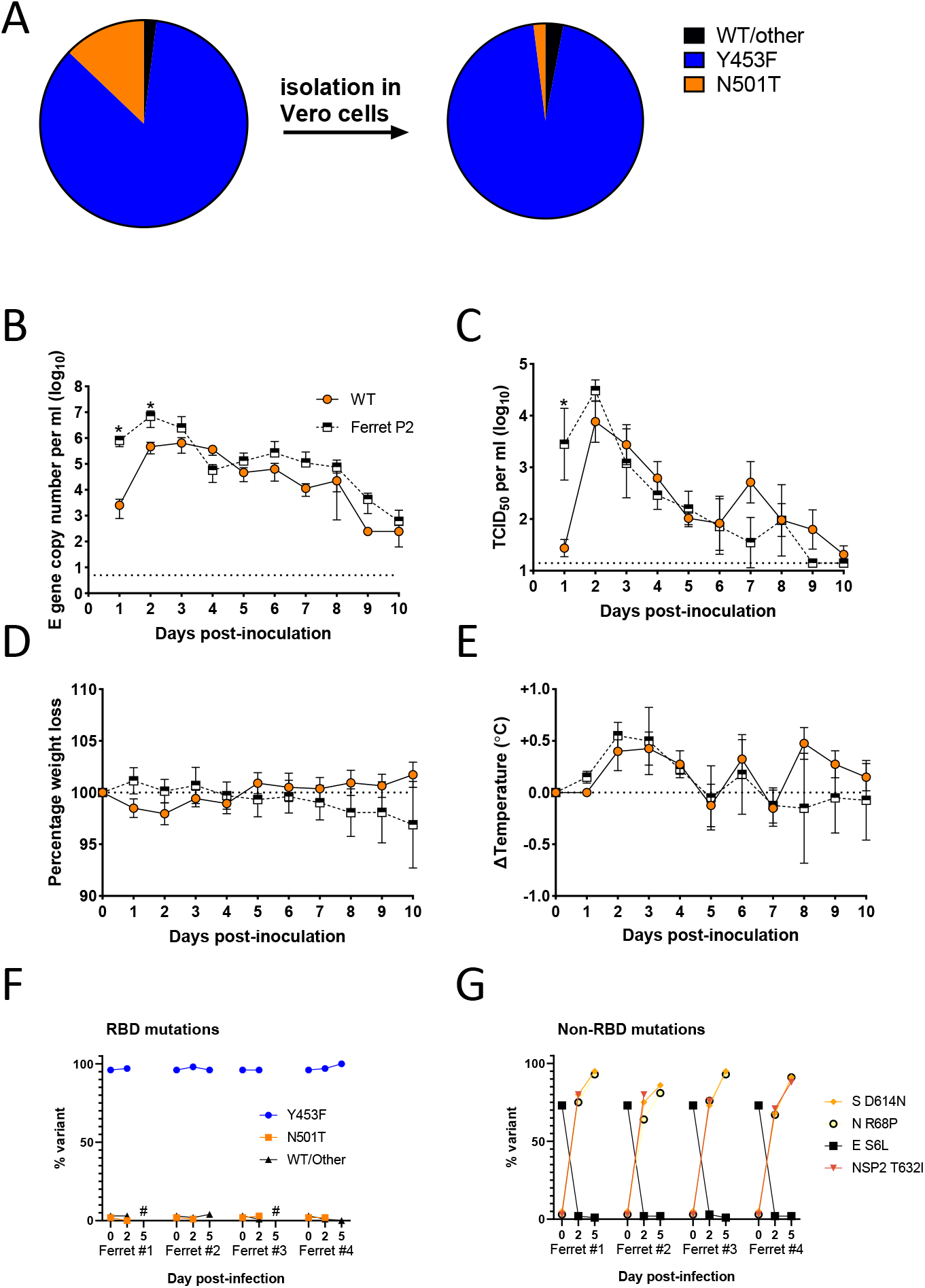
The spike mutation, Y453F, enhances replication and morbidity in ferrets. A) Deep sequencing of RBD mutations of SARS-CoV-2 from ferret passage 2 swab (see Figure 1a) before and after isolation in Vero cells. RNA (B) and infectious virus (C) shedding dynamics of ferrets directly infected with either WT (orange circles; as previously described in ^3^) or Y453F (ie ferret passage 2; black and white squares) SARS-CoV-2. N=4 naïve ferrets in each group were infected with 10^5^ p.f.u. of either virus. Percentage weight loss (D) and change in body temperature (E) were recorded daily. Statistics on B and C determined by multiple Mann-Whitney tests. *0.05 ≥ *P* Spike RBD (F) and non-RBD (G) mutations seen in Vero grown ferret passage 2 virus (time 0) from Figure 2A-D and dynamics over time in directly infected ferrets.

### Virus with Y453F shows enhanced replication and trended towards higher morbidity in ferrets

To investigate whether the Y453F-containing virus showed greater replication in ferrets, we intranasally inoculated 4 naive ferrets with ferret P2 virus and compared levels of virus shed from the nose to 4 ferrets previously inoculated with the same infectious titre of parental England/2/2020 virus ^3^(the same donors from Figure 1A). At days 1-2, the mean titre of Y453F virus shed in nasal washes was significantly higher than that of the parental virus, as determined by both E gene copy number and TCID_50_ (Figure 2B,C). Both groups of ferrets showed comparable patterns of fever during infection, peaking between days 2-4, and the Y453F-infected ferrets trended towards more weight loss over the course of the experiment (Figure 2D,E). The titre of parental virus shed and fever in parental virus-infected animals approached that in the ferret P2 infected animals by days 3-4, likely because the parental virus had gained ferret-adapting mutations, such as Y453F or N501T, by this point (see Figure 1A). Deep sequencing of the virus from the ferrets inoculated with the Y453F-containing ferret P2 virus showed the Y453F substitution was maintained in all 4 animals throughout the course of infection (Figure 2F). The E gene substitution S6L, however, was rapidly selected against, indicating that this substitution could have been an adaptation to cell culture, selected in Vero cells during isolation and amplification of the virus from nasal wash (Figure 2G). Several further substitutions, all present at very low levels in the inoculum, rapidly grew to fixation in all 4 Y453F-infected ferrets. These encoded mutations in spike at D614N, in N protein at R68P and in the NSP2 protein at T632I (Figure 2G). It is unclear whether these substitutions are all *bona fide* ferret adaptations or mutations hitchhiking as part of a selective sweep. Spike D614N may exert a similar effect to the ubiquitous SARS-CoV-2 human adaptation D614G, to non-specifically enhance ACE2 binding by promoting the spike open conformation ^27^. Overall, these data suggest that Y453F adapts the virus to ferret infection, but also further adaptations may arise during ongoing adaptation in mustelid hosts.

### Y453F enhances cell entry using the mustelid ACE2 receptor

Next, we tested whether Y453F and the other mustelid associated spike mutations improved the use of the otherwise suboptimal ferret ACE2 ^28^. We created a library of spike expression constructs, generated lentivirus-based pseudoviruses and assessed the entry of these into cells transiently expressing ACE2 from human, ferret or rat, or empty vector, as previously described ^28^. We note that ferret ACE2 differs from that of mink by only two amino acid residues that are distal to the spike interaction interface, and therefore can be considered representative for both mustelid species (See Extended Data Figure S1).

While WT (D614G) spike uses ferret ACE2 poorly for entry (>10-fold less well than human ACE2), the adaptations Y453F, N501T or F486L, as well as full Cluster 5 spike (Δ69/70, Y453F, D614G, I692V, M1229I), all allowed SARS-CoV-2 spike expressing pseudoviruses to enter into human- or ferret-, but not rat-, ACE2 expressing cells with much greater efficiency (Figure 3A, B, Extended Data Figure 2A). A nearby substitution, L452M, which has also appeared in at least one mink farm outbreaks ^11^ has no effect suggesting this is not a specific adaptation to mink (Figure 3A). Furthermore, this effect was not dependent on the presence or absence of D614G, as Y453F in a 614D background showed a similar effect (Figure 3C, Extended data Figure 2B). Consistent results were also seen using a cell-cell fusion assay (Extended data Figure 2C,D). Examining the structure of the spike RBD/ACE2 interface, each of these mink/ferret-adaptations is close to residues that differ between human and mustelid ACE2, as others have previously modelled ^29^. For example, Y453F lies close to H34Y (histidine in human ACE2, tyrosine in mustelid), N501T lies close to G354R, and F486L lies between ACE2 residues L79H, M82T and Q24L (Figure 3D).

**Figure 3.**
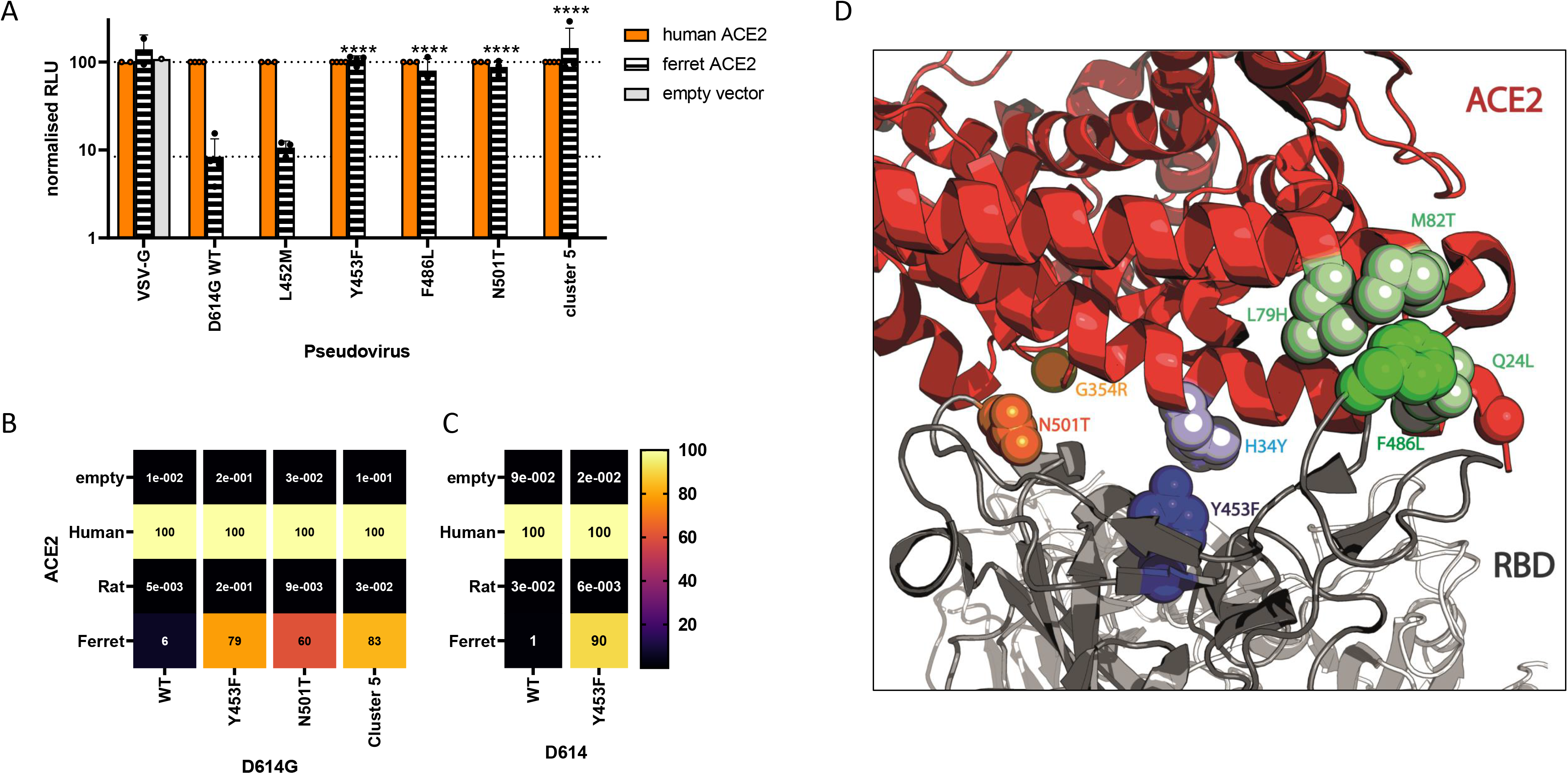
Mink- and ferret-associated spike mutations allow more efficient entry into cells expressing the ferret ACE2 receptor. A) Pseudovirus entry in human or ferret ACE2 expressing cells. Mutant SARS-CoV-2 spike-containing pseudovirus entry into HEK 293Ts expressing human or ferret ACE2 or empty vector. Entry normalised to signals from human ACE2 expressing cells. Each data point indicates mean value taken from a completely independent repeat (N≥3). Statistics were determined by comparing log-transformed values of ferret ACE2 entry using a one-way ANOVA with multiple comparisons against the WT. *0.05 ≥ *P* > 0.01; **0.01 ≥ *P* > 0.001; ***0.001 ≥ *P* > 0.0001; \*\*\*\**P* ≤ 0.0001. Entry of SARS-CoV-2 spike mutant-expressing lentiviral pseudotypes into BHK-21 cells expressing different mammalian ACE2 proteins. Pseudovirus shown contain either D614G (B) or D614 (C). Entry normalised to entry into human ACE2 expressing cells. Representative repeat shown from N≥3 repeats. D) Structure of ACE2/Spike RBD interface showing key mink-adaption residues and nearby residues that differ in mustelid and human ACE2. Figure made using PyMOL (Schrödinger) and PDB: 7A94^60^.

### Viruses containing Y453F mutation are attenuated for replication in primary human airway epithelial cells

To assess the impact of the Y453F mutation on the replication of virus in human airway epithelium, we infected primary human bronchial cells cultured at an air liquid interface with a mix of the parental and ferret P2 viruses at a low multiplicity of infection (MOI) of around 0.1 (Figure 4A). Samples were taken 24, 48 and 72 hours post-infection and analysed by deep sequencing. The WT virus significantly outcompeted Y453F, with less than ^∼^5% of reads by 48 hours post infection containing Y453F.

**Figure 4.**
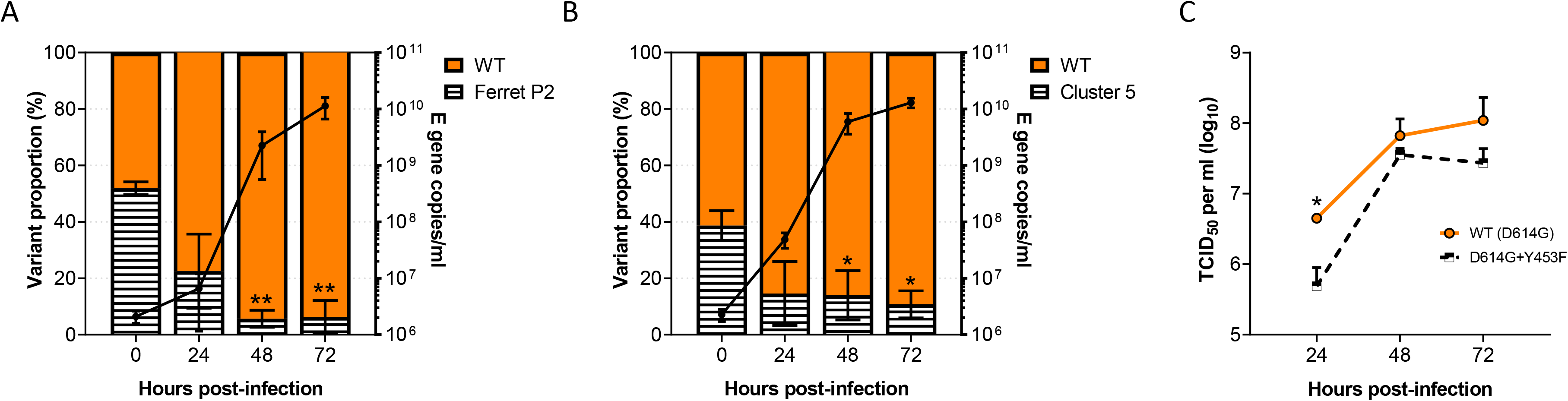
The common mink and ferret adaptation, Y453F, attenuates virus replication in primary human airway cells. Human primary airway epithelial cells cultured at air-liquid interface were infected at an MOI of approximately 0.1 with A) a mixture of parental and ferret-adapted England/2 virus B) A mixture of Mink-adapted ‘Cluster 5’ virus and a D614G control or C) either isogenic WT (D614G) or D614G + Y453F-containing reverse genetics-derived virus isolates. Virus titres were measured by TCID_50_ (C) E gene qPCR (A, B). Statistics for competition assays were determined by One-Way ANOVA with multiple comparisons against time 0. Statistics for the head-to-head growth curve (C) were determined by multiple unpaired T-tests on log-transformed data. All infections were performed on triplicate wells from matched donors (N = 3). *0.05 ≥ *P* > 0.01; \*\**P* ≤0.01.

Although the Y453F containing virus is highly similar to that which circulated in mink early in the pandemic, the most prominent zoonotic spillover from mink was the Cluster 5 virus, which further contained D614G and Δ69-70. D614G and Δ69-70 are thought to potentially enhance virus infectivity in some backgrounds ^30^. Therefore, we performed a similar competition experiment between a mixed inoculum of 40% Cluster 5 isolate and 60% early B.1 lineage, D614G containing virus (‘WT’; IC19). Again, we observed that the Y453F-containing Cluster 5 was outcompeted, constituting only ^∼^10% of reads by 24 hours post-infection (Figure 4B).

Finally, to further confirm that the attenuation of the Y453F containing viruses, particularly the ferret-adapted strain, wasn’t due to other changes in the genome (such as E S6L described above) we generated by reverse genetics (RG) two viruses on a Wuhan-hu-1, both carrying the D614G mutation in spike, WT (D614G), while the other additionally contained Y453F (D614G + Y453F). As with the ferret adapted P2 virus and Cluster 5 isolate we saw that the Y453F + D614G RG virus produced less infectious virus upon replication in the primary airway cells as compared to the otherwise isogenic WT (D614G) virus, significantly so at 24 hours post-infection (Figure 4C).

### Mink adaptation has a minimal effect on SARS-CoV-2 antigenicity

To investigate whether a mustelid-adapted SARS-CoV-2 crossing back into the human population would have a large impact on re-infections or vaccine-breakthrough we next tested whether the mutation at Y453F facilitated escape from antibody neutralization. Surprisingly, Y453F-containing ‘Ferret P2’ virus was significantly *more easily* neutralised by convalescent first wave antisera than wild type requiring only 0.6 as much antisera for a 50% neutralisation titre (Figure 5A). We further investigated the relative antigenicity of Y453F, this time using the above-described RG viruses and antisera from health care workers who had received two doses the of Pfizer-BioNtech-BNT162b2 vaccine. Again, we saw the Y453F-containing virus was more readily neutralised by 7 of the 10 vaccinee sera, although the difference was not significant (Figure 5B).

**Figure 5.**
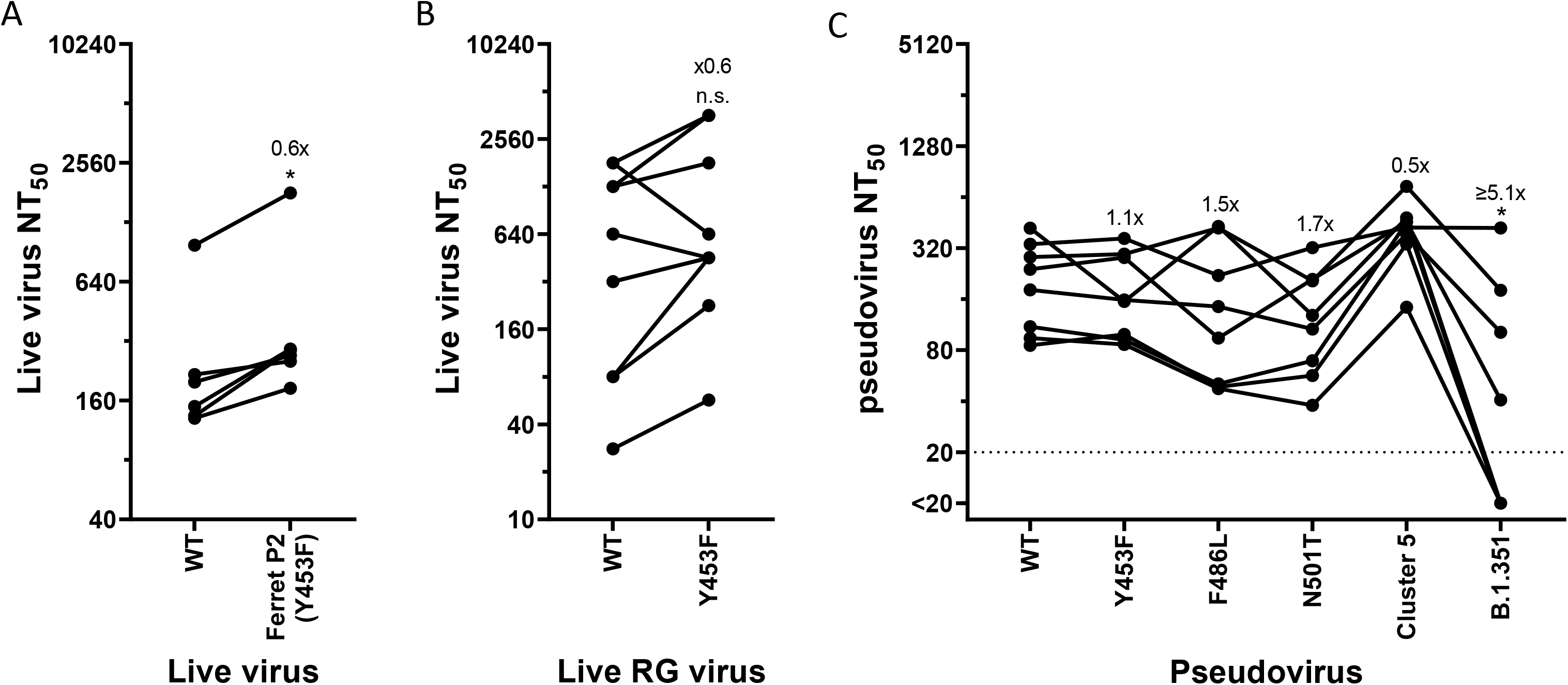
Mink and ferret associated mutations have a minimal impact on SARS-CoV-2 antigenicity. Live virus neutralisation comparing WT or Y453F-containg ferret passage 2 (A) or the isogenic reverse genetics-derived WT (D614G) and D614G + Y453F-containing SARS-CoV-2 isolates (B) using N=6 human convalescent antisera from the first UK wave (^∼^April-June 2020; A) or N=10 double-dose BNT162b2 (Pfizer-BioNTech mRNA vaccine) human antisera (B). Fold differences annotated on graph indicate differences in geometric means of NT_50_. Statistics were determined by two-tailed Wilcoxon test with matched pairs. *0.05 ≥ *P* C) Pseudovirus neutralisation of different mink-adaptations containing mutants using N=8 human convalescent antisera from the first UK wave (^∼^April-June 2020). Fold differences annotated on graph indicate differences in geometric means of NT_50_. Statistics determined by matched pair Friedman non-parametric test with multiple comparisons against WT. *0.05 ≥ *P*.

We next performed pseudovirus neutralisation assays with the previously described first wave convalescent antisera against pseudoviruses expressing the common mustelid adaptations or with full Cluster 5 spike. The B.1.351 (Beta) spike showed a significant, ^∼^5-fold drop in mean NT_50_ (Figure 5C), consistent with this virus being more difficult to neutralise with first wave antisera ^31^. None of the tested mink/ferret adaptations had any significant impact on antigenicity.

### Many circulating variants of concern show a greater ability to enter via mustelid ACE2

Following worldwide circulation of SARS-CoV-2, a number of ‘variant of concern’ and ‘variant of interest’ lineages have arisen associated with properties such as increased transmissibility, higher pathogenicity, and antigenic escape ^32^. These generally locally, or globally, outcompeted other lineages to become predominant, including the Alpha variant (B.1.1.7), first associated with infections the UK ^23^. A number of these variants have RBD mutations such as L452R, E484K and/or N501Y which are thought to promote humans ACE2 binding ^22^.

To investigate whether these variants may be more able to infect mink or ferrets than the progenitor lineage B or B.1 viruses through better use of mustelid ACE2, we again used pseudoviruses expressing these variant spike proteins and normalised entry to human ACE2 (Figure 6). We found that nearly all variants of concern tested could better utilise mink ACE2 than WT (D614G only) pseudovirus. B.1.1.7/E484K, Iota/B.1.526+E484K (first associated with infections in New York), Eta/B.1.525 (a variant with associations with West Africa) and L452R (in multiple variants of concern, including Epsilon/B.1.427/B.1.429, first associated with infections in California and Delta/B.1.617.2, which is currently replacing all other SARS-CoV-2 lineages globally) all allowed pseudovirus to utilise ferret ACE2 for cell entry to almost the same degree as human ACE2. Alpha/B.1.1.7 and Beta/B.1.351 spikes showed a much more modest boost while Gamma/P.1 (first found in Japan in travellers from Brazil) showed no improved usage of ferret ACE2. It appears L452R, E484K and N501Y may promote use of ferret ACE2, while K417N/T may result in a greater reduction in ferret ACE2 usage relative to human ACE2. Overall, these data suggest multiple circulating variants of concern may be able to infect mustelid hosts with only minimal, or indeed without, further adaptation.

**Figure 6.**
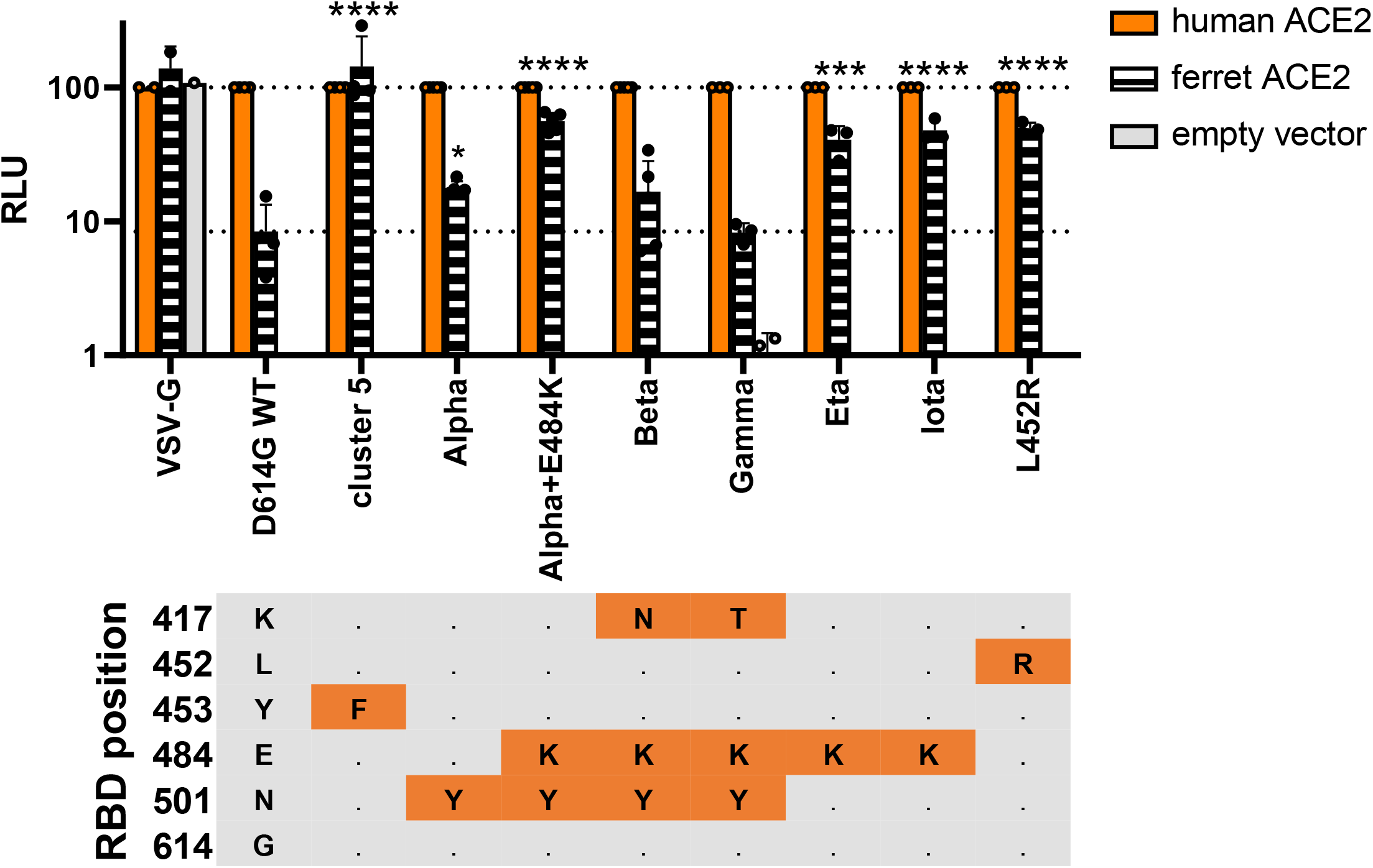
Several variants of concern show enhanced entry into ferret ACE2 expressing cells. A) Mutant SARS-CoV-2 spike-containing pseudovirus entry into HEK 293Ts expressing human or ferret ACE2, or empty vector. Entry normalised to signals from human ACE2 expressing cells. Each data point indicates data from a completely independent repeat (N≥3). Statistics were determined by comparing log-transformed values of ferret ACE2 entry using a one-way ANOVA with multiple comparisons against the WT. *0.05 ≥ *P* > 0.01; **0.01 ≥ *P* > 0.001; ***0.001 ≥ *P* > 0.0001; \*\*\*\**P* ≤ 0.0001. RBD mutational profile of the different spike proteins is shown below. Cells in orange indicate changes from WT/D614G. Alpha also known as B.1.1.7; Beta also known as B.1.351; Gamma also known as P.1; Eta also known as B.1.525; Iota also known as B.1.526+E484K.

## Discussion

In this study, we have performed a full risk assessment of mustelid hosts, such as mink and ferrets, as reservoirs for the emergence of antigenic variants or new variant of concern. We have shown SARS-CoV-2 is poorly adapted to mustelid ACE2 and therefore quickly gains adaptations, such as Y453F, N501T or F486L to utilise mustelid ACE2. However, Y453F in particular, negatively impacts replication kinetics of SARS-CoV-2 in human cells, potentially explaining why the Danish mink-origin outbreaks did not propagate further following the culling of the mink. Furthermore, in line with other studies ^16,17,31^, we found none of these mutations had a large antigenic impact, so vaccination is likely to remain effective against mustelid-adapted strains. Finally, we have shown that several VOC strains, or VOC-associated mutations, partially adapt SARS-CoV-2 spike to mustelid ACE2. Therefore, it is likely VOC lineages will continue to infect mink farms and risk spilling back over into humans.

Except for the Danish mink-adapted SARS-CoV-2 spillback, Y453F is found rarely in humans with very few isolates reported in GISAID and only a single report of the mutation arising in immunocompromised patients – this is despite Y453F having been shown in several studies to enhance human ACE2 binding, in a similar manner to the VOC-associated mutations N501Y or L452R ^22,33,34^. This would suggest that unlike the VOC-associated mutations such as N501Y, Y453F affects viral fitness in human cells. We have shown that, even in the presence of the putative stabilising NTD deletion, Δ69-70 ^30^, virus harbouring the Y453F substitution was outcompeted by a closely related virus in human cells.

Here, we have demonstrated many VOCs, particularly Alpha/B.1.1.7 as well as those containing L452R (such as Delta/B.1.617.2) could have a fundamental fitness advantage in mink by increasing interaction with mustelid ACE2, compared to previous non-variant strains. At present (August 2021), the vast majority of mink-origin SARS-CoV-2 sequences on GISAID are from the year 2020, even though there are a number of ongoing mink outbreaks reported in Europe ^35,36^, suggesting a significant reporting lag. None of the four WHO-designated variants of concern have yet been associated with mink farm outbreaks. It remains to be seen whether these VOCs would replicate in mink/ferrets without any further adaptation, but we have shown that the most common mustelid adaptations would be unlikely to have a large effect on VOC antigenicity. It will be key in the coming years to continue to closely survey farmed mink and to sequence and share any SARS-CoV-2 genomes from these animals in a timely manner as SARS-CoV-2 could still adapt in unexpected ways in mink ^37^.

This work also suggests that, particularly when investigating spike RBD mutants, ferrets (or indeed mink) are poor models for humans, as mustelid ACE2 is poorly utilised by non-adapted SARS-CoV-2 spike. Thus, it is not a given that adaptation to human ACE2 will also result in increased infectiousness, transmissibility or pathogenicity in the ferret model. However, ferrets remain a useful model for investigating non-RBD phenotypes though care should be taken to use previously ferret adapted viruses to prevent rapid adaptation.

## Methods

### Biosafety and ethics statement

All laboratory work was approved by the local genetic manipulation safety committee of Imperial College London, St. Mary’s Campus (centre number GM77), and the Health and Safety Executive of the United Kingdom, under reference CBA1.77.20.1. SARS-CoV-2 reverse genetics work was performed at CVR University of Glasgow under HSE GM notification number is GM223/20.1a. Animal research was carried out under a United Kingdom Home Office License, P48DAD9B4.

Healthcare workers convalescent antisera samples from the REACT2 studies were taken in concordance with the World Medical Association’s Declaration of Helsinki. Ethical approval was approved by the South Central-Berkshire B Research Ethic Committee (REC ref: 20/SC/0206; IRAS 283805). Sera from BNT162b2 vaccinated healthcare workers ^38^ were collected as part of a study approved by the Health Research Authority (REC ref: 20/WA/0123).

### Cells

African green monkey kidney cells (Vero; Nuvonis Technologies) were maintained in OptiPRO SFM (Life Tech) containing 2× GlutaMAX (Gibco). Human embryonic kidney cells (293T; ATCC; ATCC CRL-11268) were maintained in Dulbecco’s modified Eagle’s medium (DMEM; Gibco), 10% fetal calf serum (FCS), 1× non-essential amino acids (NEAA; Gibco), 1× penicillin-streptomycin (P/S; Gibco). Stably transduced ACE2-expressing 293T cells were produced as previously described ^3,39^, and maintained with the addition of 1 μg ml-1 puromycin to growth medium. Baby hamster kidney cells (BHK-21; ATCC CCL-10) were maintained in DMEM (Sigma-Aldrich) supplemented with 10% FCS, 1 mM sodium pyruvate solution (Sigma-Aldrich, Germany), and 1× P/S. Air–liquid interface human airway epithelium (HAEs) cells were purchased from Epithelix and maintained in Mucilair cell culture medium (Epithelix). All cell lines were maintained at 37 °C, 5% CO_2_. Cell lines were not tested for mycoplasma contamination.

### Viruses, reverse genetics and growth kinetics

The early SARS-CoV-2 strain, England/2/2020 (VE6-T) was previously isolated by Public Health England as previously described ^40^. The D614G containing strain, SARS-CoV-2/England/IC19/2020, was used as previously described ^41^. The Cluster 5 isolate - SARS-CoV-2/hu/DK/CL-5/1 – was isolated as previously described ^16^ and was kindly provided by Kevin Bewley at Public Health England. All viral stocks used in this study were grown in Vero cells in OptiPRO SFM containing 2× GlutaMAX. Virus titration was performed by median tissue culture infectious dose (TCID_50_) on Vero cells as described previously ^3^.

Virus growth kinetics and competition assays were performed as described previously ^3,42^. Briefly, in air-liquid interface HAEs, before infection cells were washed with serum-free media to remove mucus and debris. Cells were infected with 200 μL of virus-containing serum-free DMEM and incubated at 37°C for 1 h. Inoculum was then removed and cells were washed twice. Time points were taken by adding 200 μL of serum-free DMEM and incubating for 10 mins and 37°C before removal and titration.

Transformation-Associated Recombination (TAR) method in yeast was used to generate the mutant viruses described in this study. We followed essentially previously described methods ^43^ with some modifications. Briefly, a set of overlapping cDNA fragments representing the entire genome of SARS-CoV-2 Wuhan isolate (GenBank: MN908947.3) were chemically synthesized and cloned into pUC57-Kan (Bio Basic Canada Inc). Where appropriate the relevant synthetic cDNA fragment carried the mutation D614G or Y453F + D614G in the viral S gene. The cDNA fragment representing the 5’ terminus of the viral genome contained the bacteriophage T7 RNA polymerase promoter preceded by a short sequence stretch homologous to the *Xho*I-cut end of the TAR in yeast vector pEB2 ^44^. The fragment representing the 3’ terminus contained the T7 RNA polymerase termination sequences followed by a short segment homologous to the *Bam*HI-cut end of pEB2. These cDNA fragments were excised by restriction digestion and gel-extracted or PCR-amplified using appropriate primers. These fragments were then pooled and co-transformed with *Xho*I-*Bam*HI-cut pEB2 into the *Saccharomyces cerevisiae* strain TYC1 (MATa, ura3-52, leu2Δ1, cyh2^r^, containing a knockout of DNA Ligase 4) ^44^ that had been made competent for DNA uptake using the LiCl_2_-based Yeast transformation kit (YEAST1-1KT; Merck). The transformed cells were plated on minimal synthetic defined (SD) agar medium lacking uracil (Ura) but containing 0.002% (w/v) cycloheximide to prevent selection of cells carrying the empty vector. Following incubation at 30°C for 4 to 5 days, colonies of the yeast transformants were screened by PCR using specific primers to identify those carrying plasmid with fully assembled genomes. Selected positive colonies were then expanded to grow in 200 ml SD-Ura dropout medium and the plasmid extracted as described by Thao et al. (2020)^43^. Approximately 4 μg of the extracted material was then used as template to *in vitro* synthesized viral genomic RNA transcripts using the Ribomax T7 RNA transcription Kit (Promega) and Ribo m7G Cap Analogue (Promega) as per the manufacturer’s protocol. Approximately 2.5 μg of the *in vitro* synthesized RNA was used to transfect ^∼^6 ×10^5^ BHK-hACE2-N cells stably expressing the SARS-CoV-2 N and the human ACE2 genes ^45^ using the MessengerMax lipofection kit (Thermo Scientific) as per the manufacturer’s instructions. Cells were then incubated until signs of viral replication (syncytia formation) became visible (usually after 2-3 days), at which time the medium was collected (P0 stock) and used further as a source of rescued virus to infect VERO E6 cells to generate P1 and P2 stocks. Full genome sequences of viruses collected from from P0 and P1 stocks were obtained in order to confirm the presence of the desired mutations and exclude the presence of other spurious mutations. Viruses were sequenced using Oxford Nanopore as previously described ^46^.

### E gene RT-qPCR

Virus genomes were quantified by E gene RT-qPCR as previously described ^47^. Viral RNA was extracted from supernatants of swab material using the QIAsymphony DSP Virus/Pathogen Mini Kit on the QIAsymphony instrument (Qiagen). RT-qPCR was then performed using the AgPath RT-PCR (Life Technologies) kit on a QuantStudio™ 7 Flex Real-Time PCR System with the primers specific for SARS-CoV-2 E gene ^48^. For absolutely quantification of E gene RNA copies, a standard curve was generated using dilutions viral RNA of known copy number. E gene copies per ml of original virus supernatant were then calculated using this standard curve.

### Live virus neutralisation

Convalescent antisera from health care workers who had tested positive by RT-qPCR were taken from the REACT2 study as described previously ^49,50^. Double dose BNT162b2 (Pfizer-BioNtech) antisera from health care workers was generated as previously described ^38^.

Live virus neutralisation assays were performed in Vero cells as described elsewhere ^42^. Briefly serial dilutions of sera were incubated with 100 TCID_50_ of virus for 1 h at room temperature then transferred to 96 well plates of Vero cells. Plates were incubated at 37°C, 5% CO_2_ for 42 h before fixing cells in 4% paraformaldehyde (PFA). Cells were permeabilised in methanol 0.6% H_2_O_2_ and stained for 1 h with an antibody against SARS-CoV-2 nucleocapsid protein (Sino Biological; 40143-R019, 1:300 dilution). Cells were further stained with the secondary antibody anti-rabbit HRP conjugate (Sigma; 1:3000 dilution) for 1 h. TMB substrate (Europa Bioproducts) was added and developed for 20 mins before halting the reaction with 1M HCl. Plates were read at 450nm and 620nm and the concentration of serum needed to reduce virus signal by 50% was calculated to give NT50 values.

For the CPE-based neutralisation assay (reverse genetics virus vs Pfizer antisera), serial dilutions of sera were incubated with 100 TCID_50_ of virus for 1 h at 37°C, 5% CO2 in 96 well plates before a suspension of Vero-ACE2-TMPRSS2 cells were added and incubated for 3 days at 37°C, 5% CO_2_. Wells were stained using crystal violet, scored for the presence of virus-induced cytopathic effect and the reciprocal of the highest serum dilution at which protection was seen was calculated as the serum titre.

### Plasmids and cloning

Lentiviral packaging constructs pCSLW and pCAGGs-GAGPOL were made as previously described. Mutant SARS-CoV-2 expression plasmids were generated by site-directed mutagenesis using the QuikChange Lightning Multi Site-Directed Mutagenesis Kit (Agilent). Unless otherwise stated all SARS-CoV-2 spike expression plasmids were based on the Wuhan-hu-1 reference sequence ^41^, with the additional substitutions D614G and K1255*STOP (aka the Δ19 mutation or cytoplasmic tail truncation). Animal ACE2 proteins in pDisplay were generated and used as previously described ^5^.

### Pseudovirus assays

SARS-CoV-2 spike-bearing lentiviral pseudotypes (PV) were generated as described previously ^3,28^. At ICL, 100 mm dishes of 293Ts were transfected using lipofectamine 3000 (Thermo) with a mixture of pCSFLW, pCAGGS-GAGPOL and spike proteins expressed in pcDNA3.1. After 24 h supernatant was discarded and replaced. Pseudovirus-containing supernatant was collected and pooled at 48 and 72 h post-transfection, passed through a 0.45 μm filter, aliquoted and frozen at −80°C. At the Pirbright Institute pseudovirus was generated in 6-well plates. Cells were transfected using polyethyleneimine (PEI) with a mixture of pCSFLW, p8.91 and SARS-CoV-2 spikes expressed in pcDNA3.1. As before supernatant was discarded and replaced at 24 h post-transfection then harvested and pooled at 48 and 72h. Supernatant was clarified by low-speed centrifugation, aliquoted and frozen at −80°C.

Pseudovirus assays at ICL were performed as previously described ^3^. Briefly 10mm diameter dishes of 293T cells were transfected with 1 μg of ACE2 of empty vector using lipofectamine 3000. 24 h later cells media was replaced, and cells were resuspended by scraping and plated into 96 well plates and overlayed with pseudovirus. 48 h later cells were lysed with reporter lysis buffer (Promega) and assays were read on a FLUOstar Omega plate reader (BMF Labtech) using the Luciferase Assay System (Promega).

At Pirbright assays were performed largely as previously described ^28^. Briefly, BHK-21 cells were transfected with 500 ng of ACE2 or empty vector (pDISPLAY) using TransIT-X2 (Mirus Bio) according to the manufacturer’s recommendation. 24 h later, media was removed, and cells were harvested following the addition of 2mM EDTA in PBS, resuspended in DMEM and plated into white-bottomed 96 wells plates (Corning). Cell were overlayed with pseudovirus 24 h later and incubated for 48 h. Firefly luciferase was quantified whereby media was replaced with 50 μL Bright-Glo substrate (Promega) diluted 1:2 with PBS and read on a GloMax Multi+ Detection System (Promega). CSV files were exported onto a USB flash drive for analysis.

Pseudovirus neutralisation assays were performed by incubating serial dilutions of heat-inactivated human convalescent antisera with a set amount of pseudovirus. Antisera/pseudovirus mix was then incubated at 37°C for 1 h then overlayed into 96 well plates of 293T-ACE2 cells. Assays were then lysed and read as described above.

### Cell-cell fusion assay

Cell-cell fusion assays were performed as described elsewhere ^28,51^. Briefly, 293Ts stably expressing rLuc-GFP 1-7 effector cells ^52^ were transfected with empty vector, WT or mutant SARS-CoV-2 spike proteins. BHK-21 target cells stably expressing rLuC-GFP-8-11 (target cells) were co-transfected with ACE2 expression constructs. Target cells were co-cultured with effector cells 24 h post-transfection and quantification of cell-cell fusion was performed 24 h later with the *Renilla* luciferase substrate, Coelenterazine-H (Promega). Luminescence was read on a Glomax Multi+ Detection System (Promega). CSV files were exported on a USB flash drive for analysis.

### Ferret infection study

Ferret (*Mustela putorius furo*) infection studies with SARS-CoV-2 virus were performed as described previously ^3^. All ferret studies were performed in a containment level 3 laboratory, using a bespoke isolator system (Bell Isolation Systems). Outbred female ferrets (16–20 weeks old) weighing 750–1,000 g were used. Four donor ferrets were inoculated intranasally with 200 μl of 10^5^ p.f.u. of each virus while lightly anaesthetized with ketamine (22 mg kg−1) and xylazine (0.9 mg kg−1).

Prior to the start of the study ferrets were confirmed to be seronegative to SARS-CoV-2. All animals were nasal-washed daily, while conscious, by instilling 2 ml of PBS into the nostrils; the expectorate was collected into disposable 250-ml sample pots. Ferrets were weighed daily post infection, and body temperature was measured daily via subcutaneous IPTT-300 transponder (Plexx B.V).

### RNA extraction and sequencing

For Sanger sequencing, RNA was extracted from nasal washes using QIAamp viral RNA mini kit (Qiagen). RNA was reverse transcribed using Superscript IV (Invitrogen) and PCR of the spike was performed using KOD polymerase (Merck). For next generation sequencing RNA from virus-containing samples were extracted using the QIAsymphony DSP Virus/Pathogen mini kit (Qiagen). RNA was DNase-treated using the TURBO-free Kit (Invitrogen; (AM1907). cDNA was synthesised using the superscript IV reverse transcriptase (Invitrogen) and random primer mix (NEB) before amplification by the ARTIC Network protocol using the multiplexed primer scheme version 3. Fast5 files were basecalled with guppy (v.5.0.7) with high accuracy calling (hac). The fastq files produced by Nanopore sequencing were filtered with lengths 400 and 700 using Artic-ncov2019 pipeline v1.2.1 (https://artic.network/ncov-2019/ncov2019-bioinformatics-sop.htμl) by “artic guppyplex” function. The other function of “artic minion” in the Artic-ncov2019 pipeline with “--medaka --medaka-model r941_min_high_g360 --normalise 0” parameters was then used to process the filtered fastq files to generate ARTIC V3 primer trimmed bam files and consensus genome sequences. These primer trimmed bam files were further analysed using DiversiTools (http://josephhughes.github.io/btctools/) with the “-orfs” function to generate the ratio of amino acid change in the reads and coverage at each site of protein in comparison to the reference SARS-CoV-2 genome (MN908947.3) as we previous description ^53^.

### Phylogenetic analysis

All sequences with host species labelled as *Neovison vison* were retrieved from the Global Initiative on Sharing All Influenza Data (GISAID) database (sequences retrieved on 7 July 2021). A table of accession IDs and acknowledgement is given in Supplementary Table S1. A sequence with only 397 nucleotides (hCoV-19/mink/Spain/NV-2105/2021, EPI_ISL_1490748) was excluded from analysis. Sequences were aligned to the Wuhan-Hu-1 reference genome sequence (MN908947) ^54^ using MAFFT v7.475 ^55^ and the alignment was then checked manually. Seven further sequences were excluded from further analysis as they lacked nucleotide data enabling the determination of amino acid identity at spike positions 453, 486, 501 or 614 (hCoV-19/mink/USA/MI-CDC-II1O-7265/2020, EPI_ISL_925307; hCoV-19/mink/USA/MI-CDC-IHWB-7153/2020, EPI_ISL_925308; hCoV-19/mink/USA/WI-CDC-CX2X-2436/2020, EPI_ISL_1014948; hCoV-19/mink/Netherlands/NB-EMC-3-5/2020, EPI_ISL_523094; hCoV-19/mink/Netherlands/NB-EMC-3-4/2020, EPI_ISL_523093; hCoV-19/mink/Netherlands/NB-EMC-40-4/2020, EPI_ISL_577788; hCoV-19/mink/Denmark/mDK-56/2020, EPI_ISL_641448). Epidemiological lineages were determined using the Pangolin COVID-19 Lineage Assigner ^56,57^ (pangolin v3.1.5, pangoLEARN v15/06/2021). Phylogenetic analysis was performed using the remaining 936 mink genomes rooted on the Wuhan-Hu-1 reference genome (MN908947) with a general time reversible model of nucleotide substitution, a proportion of invariant sites estimated from the data and a gamma distribution describing among-site rate variation (GTR + / + Γ) built using RAxML v8.0.0 ^58^ with the phylogeny rooted on the sequence of the virus Wuhan-Hu-1. The maximum likelihood phylogeny was plotted, alongside data on sampling location extracted from the virus name and amino acid identity at spike positions 453, 486, 501 and 614, in R using the *ggtree* package ^59^.

### Statistics and reproducibility

Statistics throughout this study were performed using one-way analysis of variance (ANOVA) or Student’s *t*-test and are described in the figure legends. No statistical method was used to predetermine sample size. Several genome sequences were manually removed from the phylogenetic analysis and were described in the associated sections. The experiments were not randomized, and the investigators were not blinded to allocation during experiments and outcome assessment.

## Acknowledgements

The SARS-CoV-2 virus isolate, England/2 was provided by Public Health England, and we thank M. Zambon, R. Gopal and M. Patel for their help. The authors would also like to thank Kevin Bewley of Public Health England for help obtaining the Cluster 5 isolate, SARS-CoV-2/hu/DK/CL-5/1 and Michelle Willicombe, Maria Prendecki and Candice Clarke for their help obtaining the Pfizer double dose antisera. We also thank E. J. Louis, University of Leicester for generously providing the TAR in yeast system. The authors thank all researchers who have shared genome data openly via the Global Initiative on Sharing All Influenza Data (GIASID).

This work was supported by the G2P-UK National Virology Consortium funded by the MRC (MR/W005611/1). Additional funding to DB, AM, NT and GG were funded by The Pirbright Institute’s BBSRC institute strategic programme grant (BBS/E/I/00007038). SARS-CoV-2 research for JAH, RPR, HG, ID-B, XD and NPR is supported by the U.S. Food and Drug Administration Medical Countermeasures Initiative contract (5F40120C00085). The work at the CVR was also supported by the MRC grants (MC_UU12014/2) and the Wellcome Trust (206369/Z/17/Z).

The article reflects the views of the authors and does not represent the views or policies of the FDA.

## Contributions

JZ, TPP, JB, DHG, DB and WSB conceived and planned the experiments. JZ, TPP, JCB, DHG, AMEE, RP-R, VMC, GDL, WF, WTH, RK, LB, RF, RL, NT, GG, HG, ID-B, XD, NPR, FS, MCG and PFM performed the experiments and analysed the data. AHP, MP, JAH, DB and WSB provided supervision. TPP and WSB wrote the manuscript with input from all other authors.

## Extended data Figure legends

**Extended data Figure 1.**
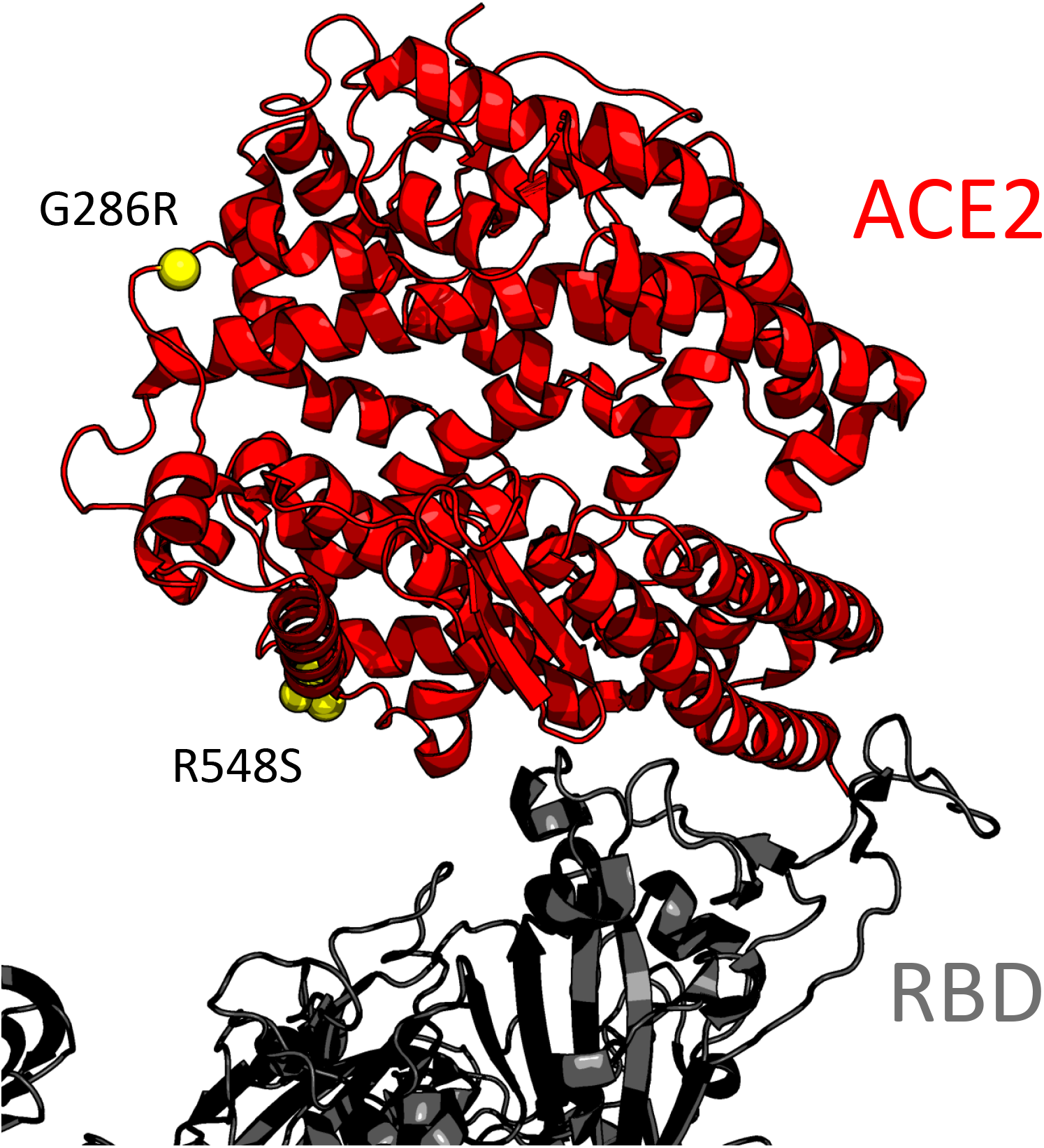
Amino acid differences between ferret and mink ACE2. Differences between ferret and mink ACE2 are shown on the structure of human ACE/Spike structure PDB: 7A94^60^.

**Extended data Figure 2.**
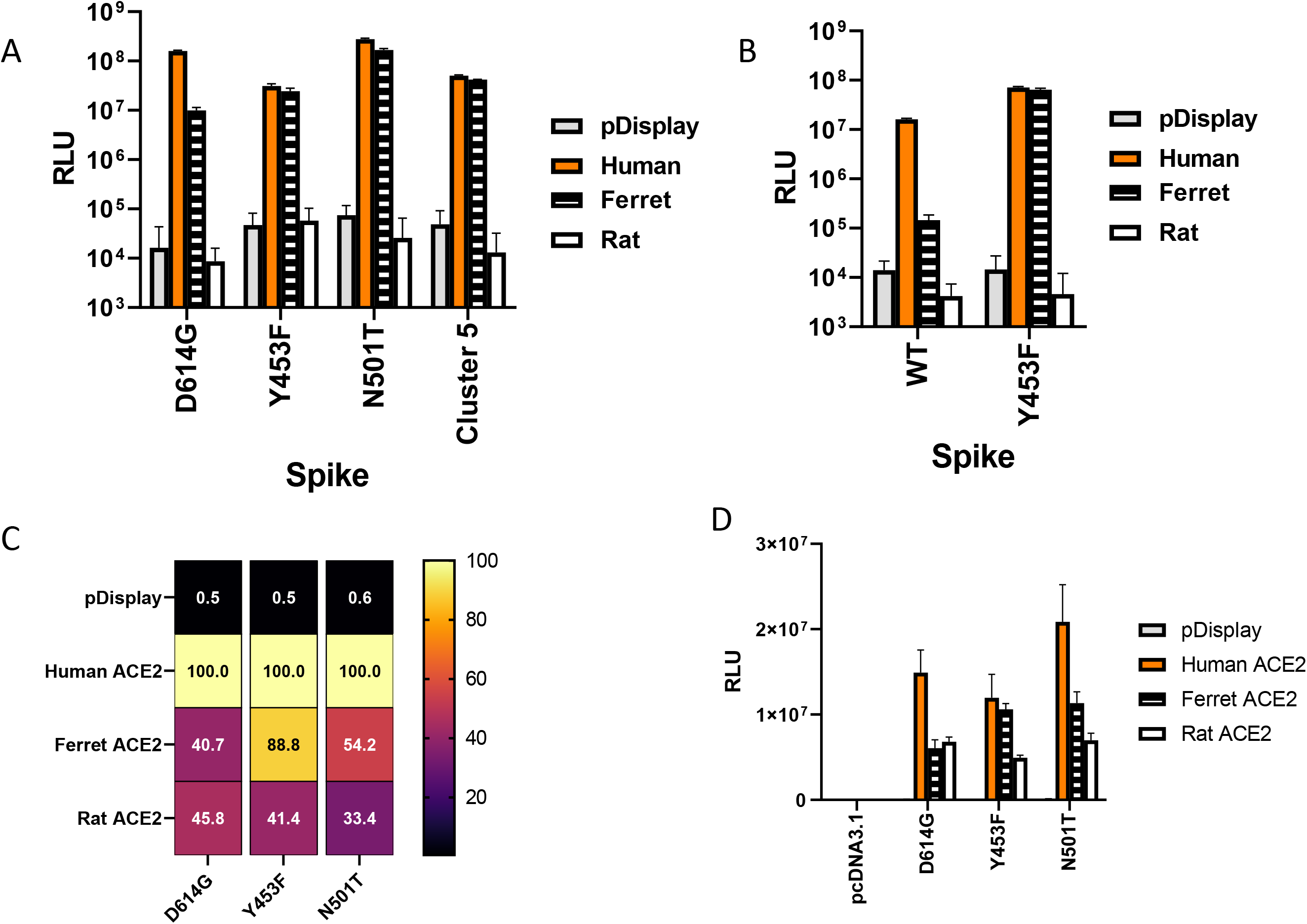
Extended data from Figure 3. A) Non-normalised data from Figure 3B. B) Non-normalised data from extended data Figure 3C. C, D) Cell-cell fusion assays of HEK 293Ts with rLUC-GFP1-7 transfected with the stated Spike protein and BHK-21 cells expressing the named ACE2 and rLUC-GFP 8-11. Luminescence values shown normalised to human ACE2 (C) or as raw values (D).

